# Identification of genes promoting fitness of a plant-associated *Salmonella* Choleraesuis strain on alfalfa sprouts during cold storage

**DOI:** 10.64898/2026.01.24.701464

**Authors:** Maximilian Beck, Laura Führer, Steffen Porwollik, Weiping Chu, Verena Hohenester, Irmak Sah, Michael McClelland, Claudia Guldimann, Irene Esteban-Cuesta

**Affiliations:** Competence Center for Food Safety, Chair of Food Safety and Analytics, Veterinary Faculty, LMU Munich, Oberschleissheim, Germany; Department of Microbiology and Molecular Genetics, School of Medicine, University of California, Irvine, California, United States of America

**Keywords:** transposon insertion sequencing, salmonellosis, microbial contamination, transposon library, food safety

## Abstract

Consumption of sprouted seeds, such as alfalfa sprouts, has increased in recent years due to their perceived health benefits. However, these food products have repeatedly been associated with outbreaks of foodborne pathogens, including *Salmonella enterica* serovars. An *S. enterica* serovar Choleraesuis strain previously isolated from melon fruit internal tissues was selected as a model to explore plant-pathogen interactions on alfalfa sprouts. Using this strain, we generated a barcoded transposon mutant library comprising approximately 33,000 unique insertions. This library and a collection of individual insertion mutants derived from it were used to identify genetic mechanisms contributing to the fitness of this *S.* Choleraesuis strain on alfalfa sprouts. The library was screened on sprouts during cold storage at 8°C. Negative selection for mutants with insertions in *eda, fabF, lpp1_2, pnp, stpA, SCHChr_03621* and two intergenic regions were identified. Competition experiments between individual insertion mutants and the wild type confirmed the phenotype of three genes: *eda*, coding for a keto-hydroxyglutarate-aldolase/keto-deoxy-phosphogluconate aldolase involved in the Entner-Doudoroff pathway, *mnmG*, encoding the glucose-inhibited division protein, and *fabF*, involved in fatty acid biogenesis. This study offers a genome-wide perspective on the genes enabling a plant-associated *Salmonella* strain to persist on alfalfa sprouts. We highlight factors that are critical not only for persistence throughout the entire cold-storage period under conditions that closely simulate real shelf-life conditions in this high-risk food matrix.

## 1 Introduction

Sprouts are germinated seeds that are harvested before true leaves develop and generally consumed raw (Official Journal of the European Union, 2017). Due to their high nutrient concentration and antioxidant content, sprouts are perceived as healthy foods, and their consumption has increased (Benincasa et al., 2019). However, the warm, humid production environment in which sprouts are produced is ideal for microbial proliferation. As a result, sprouts are frequently associated with elevated microbial loads, including a notable prevalence of antimicrobial-resistant bacteria (Rahman et al., 2022; Young Kim et al., 2022).

Multiple foodborne outbreaks have been linked to contaminated sprouted seeds, specifically involving *Escherichia coli* O157:H7 (Frank et al., 2011) and several *Salmonella enterica* serovars (Ma et al., 2024). *S. enterica* is of particular concern due to its significant global public health impact (EFSA & ECDC, 2025) and frequent involvement in sprout-associated outbreaks (Caulcrick-Grimes et al., 2026; EFSA, 2025; Miyahira & Antunes, 2021). This highlights the ability of *S. enterica* to persist throughout the entire sprout production and distribution system, ultimately reaching consumers.

Previous studies have explored the mechanisms of *S. enterica* attachment to sprouts during the germination process at room temperature (Barak et al., 2005; Holden et al., 2024) as well as the interactions of two epidemiologically dominant serovars, Typhimurium and Enteritidis, under conditions that mimic the shelf-life of sprouted seeds under refrigeration (Führer et al., 2025). However, less is known about the adaptive responses of other serovars, particularly strains isolated from plant-associated sources.

In this study, we investigate the genetic determinants required for the fitness of an *S. enterica* serovar Choleraesuis isolate during cold storage on alfalfa sprouts. This strain was previously shown to internalize into muskmelons (Esteban-Cuesta et al., 2018), suggesting a tolerance of plant-associated environments. To explore this, we generated a barcoded transposon insertion sequencing (TIS) library pool as well as an ordered library of individual insertion mutants in this *S. enterica* serovar Choleraesuis strain, 11-Ga-2015 (SCH).

Overall, this study examines genome-wide fitness mechanisms in a serovar traditionally considered host-adapted to pigs - a group rarely implicated in human infection yet often associated with increased disease severity when infections occur (Jones et al., 2008). The ability of this strain to persist on a plant substrate under cold storage conditions highlights the potential for cross-kingdom adaptation within this important pathogen.

## 2 Material and Methods

### 2.1 Barcoded Transposon Insertion Sequencing library construction in *S.* Choleraesuis

*Salmonella enterica* subsp. *enterica* serovar Choleraesuis strain 11-Galia-2015 (SCH) (Esteban-Cuesta et al., 2021) was used to construct a barcoded TIS library. This strain was originally isolated from the pulp of a Galia melon (Esteban-Cuesta et al., 2018) from a German supermarket in 2015, and had been preserved at −80°C in Luria-Bertani broth (LB; Carl Roth GmbH & Co. KG, Karlsruhe, Germany) with 20% glycerol (Th. Geyer GmbH & Co. KG, Renningen, Germany).

The construction of the barcoded Tn*5*-based mutant library was performed as previously described (de Moraes et al., 2017; de Moraes et al., 2018), with minor modifications. Briefly, an N_21_ random barcode flanked by Illumina primer sequences was inserted into the EZ-Tn*5* transposon via a two-step PCR, using primers Tn*5*PCR1L and Tn5PCR1R in the first step and the following cycling conditions: 98°C 90 s; 35 × (98°C 10 s, 62°C 20 s, 72°C 115 s); final extension 72°C 3 min; hold 8°C. The second PCR step was performed using primers 5P-Tn5PCR2L and P-Tn5PCR2R with similar cycling conditions: 98°C 90 s; 30× (98°C 10 s, 61°C 20 s, 72°C 115 s); final extension 72°C 3 min; hold 8°C. All primer sequences are shown in **Supplementary Table S1.** The purified construct was subsequently randomly introduced into the SCH genome using EZ Tn*5* transposase (Epicentre Biotechnologies, Madison, WI, USA), as per the manufacturer’s recommendations. Transformed cells were harvested from LB^kan60^ (60 µg/ml kanamycin, Carl Roth) agar after an overnight culture at 37°C.

The methods for mapping the barcoded transposons to specific locations in the genome have previously been described (de Moraes et al., 2017). For the barcoded SCH TIS library, extraction of the genomic DNA was performed using the GenElute bacterial genomic DNA kit (Sigma-Aldrich, St. Louis, Missouri, USA), followed by a standard 2h linear amplification reaction with a temperature ramp to 37°C using a random N_10_ primer (KlenowN_10_) and exo-Klenow enzyme (New England Biolabs, Ipswich, MA, USA; NEB). Deactivation was performed at 75°C for 20 min, and products were purified using the QIAquick PCR Product Purification kit (Qiagen, Maryland, USA). The region that includes the N_21_ barcode and the genomic DNA adjacent to the transposon was amplified using a nested PCR protocol that included Illumina primer sequences. Primer pair Klenow_PCR1SCHL/Klenow_PCR1SCHR was used in the first step, whereas Klenow_PCR2SCHL, including the Illumina P5 adapter, was paired with outward-facing indexed primers that included the Illumina Read2 and P7 adapter in the second step. After QIAquick purification of the resulting amplicons, paired-end 150-base reads were obtained, and the relevant 71 base-region was mapped to the *de novo* assembled SCH genome using CLC Genomics Workbench v24 (Qiagen, Venlo, Netherlands). Approximately 33,000 individual barcodes were mapped to the SCH genome.

### 2.2 Establishment of the ordered SCH transposon library

To make an ordered SCH transposon library, 9,595 individual SCH Tn mutant clones were toothpicked from LB^kan60^ agar plates during the generation of the TIS library prior to pooling. They were grown separately in LB^kan60^ in a total of 101 96-well plates (“source plates”). All mutants from each plate were pooled (resulting in 101 “plate pools”, each consisting of 95 mutants), and all mutants from the same wells across all plates were also pooled (resulting in 95 “well pools”, each with 101 mutants). The 101 “plate pools” and the 95 “well pools” were then prepared for sequencing by proteinase K digestion and PCR amplification using primers PCRSCHoTnL and PCRSCHoTnR to create Illumina sequencing libraries. Illumina sequencing was used to identify each N_21_ barcode. Analysis of barcode occurrence in the “plate pools” and “well pools” revealed the exact plate/well coordinates of each barcoded mutant in the “source plates”, while prior mapping had revealed the location of each barcoded transposon insertion in the SCH genome.

For the final selection, each mutant that was the sole representative of a genetic feature of SCH was included. When multiple mutants were available for a given gene, two clones per gene were selected. Priority was given to clones with transposon insertions on opposite strands of the gene, preferably near the N-terminus, while maintaining a minimum distance of 10 bases from gene boundaries.

### 2.3 Comparative growth analysis

Population dynamics of the barcoded SCH TIS library on alfalfa sprouts were performed in five replicates and statistically compared to those of the wild type strain 11-Galia-2015 (WT). Inoculation and sampling were performed at 8°C as described for the library screenings below. An ANOVA test was used for statistical analysis, with adjusted *p*-values<0.05 considered statistically significant.

### 2.4 Library screening on alfalfa sprouts during cold storage

The methods for the TIS library screen on food matrices were previously described (Esteban-Cuesta et al., 2026; Führer et al., 2025). In brief, 300 µl of library stocks were thawed and propagated in 30 ml of LB^kan60^ broth at 37°C and 200 rpm until OD_600_ reached 1.0. After centrifugation of the 10-fold diluted culture (4,500 rcf, 5 min), the supernatant was discarded, and the inoculum was prepared by resuspending the cell pellet in 5 mL of phosphate-buffered saline (PBS; Carl Roth).

Alfalfa sprouts (*Medicago sativa* L.) were purchased from local supermarkets. To quantify the background microbiota of the uninoculated sprouts, mesophilic aerobic bacteria (MAB) were enumerated according to EN ISO 4833-2:2013, and *Enterobacteriaceae* were quantified following EN ISO 21528-2:2017 (anaerobic incubation conditions: O_2_ <0.1%, CO_2_ 7.0–15.0%) at each sampling time point. The absence of *Salmonella* spp. in sprout samples was verified by qualitative microbiological analysis, in accordance with EN ISO 6579-1:2017.

For inoculation, 10 g of alfalfa sprouts were aseptically transferred into sterile filter Stomacher^®^ bags (0.4 L, Avantor VWR International GmbH, Darmstadt, Germany). Each sprout sample was mixed with 5 ml of the inoculum, while control samples received only 5 ml of PBS. The final inoculum concentration was approximately 8.0 log₁₀ CFU/ml with a final concentration of 7.0 log₁₀ CFU/g sprouts to maintain library complexity. All samples were incubated at 8°C for 5 days.

Sampling of the mutant libraries was performed directly from the inoculum as well as at 1h (d_1_), 48h (d_3_), and 96h (d_5_) post-inoculation. At these time points, alfalfa sprouts and negative control samples were transferred to 90 mL of LB^kan60^ in Erlenmeyer flasks and incubated at 37°C with shaking at 200 rpm for 30 min to detach bacteria from the matrix. *Salmonella* populations were subsequently quantified by plating serial dilutions on BPLS Agar (Merck KGaA, Germany), which was incubated at 37°C for 24 h. Population dynamics of the barcoded SCH TIS library on sprouts and in PBS were statistically compared using ANOVA, with *p*_adj_-values<0.05 considered significant.

In parallel, samples in 90 ml LB^kan60^ were filtered through the stomacher bags and centrifuged (4.500 rcf, 5 min) to eliminate residual plant material in the medium, and the pellet was resuspended in 90 ml fresh LB^kan60^. Sprouts samples and negative controls were then further incubated for 7.5h at 37°C and 200 rpm. From this, 1 ml aliquots were stored at −80°C in broth without glycerol for subsequent genomic DNA extraction and sequencing. Each experiment was performed in five biological replicates.

For sequencing library preparation, 40 μl aliquots were washed three times with sterile ultrapure water and pelleted. 15 µl washed cells were subjected to a two-steps lysis protocol. In the first step, cells were incubated with 1.6 µl lysozyme (10 mg/ml; AppliChem GmbH, Darmstadt, Germany) in 8 µl lysis buffer consisting of 10 mM Tris [pH 8.0], 1 mM EDTA (Carl Roth), and 0.1% Triton X-100 (Carl Roth) for 5 min at 20°C, followed by enzyme inactivation at 98°C for 5 min and cooling to 4°C. In the second, 1 µl proteinase K (100 mg/ml; Sigma-Aldrich) was added in 5 μl lysis buffer, incubated for 2h at 55°C with subsequent proteinase K inactivation at 95°C for 10 min, and immediate cooling. After centrifugation for 1 min at 14,100 rcf, 5 μl of the lysate were used as template for PCR amplification with indexing primers annealing to sequences flanking the transposon barcode (Illumina-barcoded primers N and S; 0.2 μM each; see Table S1). PCR amplification was carried out using Q5™ Hot Start High-Fidelity 2× Master Mix (NEB) at a final 1× concentration. The cycling protocol consisted of an initial denaturation at 98°C for 2 min, followed by 10 cycles of 98°C for 10 s, 65°C for 10 s, and 72°C for 20 s. Additional 20 cycles were then performed with denaturation at 98°C for 10 s and extension at 72°C for 20 s. A final elongation step was conducted at 72°C for 2 min, after which reactions were held at 20°C. Amplified products were pooled in equimolar volumes and purified using the QIAquick PCR Purification Kit (Qiagen) according to the manufacturer’s protocol. Sequencing libraries were analyzed by dual-indexed paired-end sequencing (100-150 bp reads) on an Illumina NovaSeq 6000 platform. On average, 1,057,353 mapped barcodes (a minimum of 736,054 and a maximum of 1,417,144 mapped barcodes) were sequenced per sample.

### 2.5 Bioinformatic analysis

Raw reads from different samples were demultiplexed by Illumina index and analyzed with custom Python scripts to identify and quantify barcodes flanked by invariant sequences. Barcode counts were aggregated per disrupted gene, and changes in mutant abundance across time points and between matrix and *no matrix* samples were statistically assessed using DESeq2 (Love et al., 2014), with log_2_-fold changes (fc) reported **(Table S2)**.

Mutations were considered to have a significant fitness effect for SCH on alfalfa sprouts if, at a given sampling time point, it showed: (i) a log_2_-fc>|1.0| with a *p*_adj_*-*value ≤0.01 when comparing the populations on sprouts versus PBS (*no matrix* sample), and (ii) the same significance threshold (log_2_-fc >|1.0|, adjusted *p*_adj_ ≤ 0.01) when comparing the inoculum to the sprout sample at the same sampling time point. This dual-filtering approach ensured that only the genes most relevant to the interaction between SCH and sprouts were identified.

To prepare gene lists for input into KEGG (Kanehisa et al., 2023) and Gene Ontology (GO) (Aleksander et al., 2023; Ashburner et al., 2000; Thomas et al., 2022) enrichment analyses, the adjusted *p*-value threshold for significance was relaxed from ≤0.01 to <0.05. GO enrichment was performed using the TopGO package (Alexa & Rahnenfuhrer, 2024), while KEGG pathway analysis was conducted with the enrichKEGG function from the ClusterProfiler package (Wu et al., 2021) in R (v4.4.1). KEGG pathways with *q*-values ≤0.01 and GO terms with classic Fisher *p*-values ≤0.01 were considered enriched.

### 2.6 Competition assays

The phenotypes of candidate genes identified in the transposon analysis were verified using selected individual insertion mutants (IM) from the ordered SCH transposon library in direct competition with the WT strain. For some of these candidate genes, two mutants were available – one harboring the transposon insertion in the sense direction and the other harboring the transposon insertion in the antisense direction – facilitating the identification of orientation-dependent effects. The insertion mutants analyzed in this study are listed in **Table 3**.

For competition assays, a single colony of the IMs and the WT strain were cultured overnight at 37°C with shaking at 200 rpm in 5 ml LB^kan80^ (80 µg/ml) and LB, respectively. From the overnight cultures, 300 µl were cultured in 30 ml LB^kan80^ (80 µg/ml) and LB, respectively, under the same conditions until OD_600_ 0.9 was reached. IM and WT pairs were then mixed at a 1:1 ratio, diluted 1:10 in PBS, and 1 ml of this IM-WT mix was centrifuged. Pellets were resuspended in 5 ml PBS, and the centrifugation and resuspension steps were repeated once to wash the cells. Subsequently, 5 ml were used to inoculate 10 g sprout samples and corresponding no matrix control bags containing only the inoculum mixture. To control nutrient-dependent effects, parallel competitions were resuspended in LB medium instead of PBS. The final inoculum concentration on the samples was approximately 7.0 log_10_ CFU/ml or g sprouts. All samples were incubated at 8°C. The specific incubation period for each candidate gene was selected based on the time point showing the strongest fitness effect in the transposon screen. To assess changes in population ratios, samples were subsequently plated on both BPLS and BPLS^kan80^ agar and incubated at 37°C for 24 h. WT colony counts were calculated by deducting the CFUs observed on BPLS^kan80^ from those on BPLS.

The phenotype of the candidate genes was confirmed by comparing colony count changes of the respective sense and antisense IM and the WT strain, and calculating the corresponding competition indices 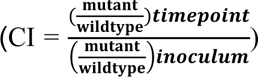 for each experiment. A negative effect was confirmed if the average CI of all the replicates plus the standard deviation was <1. If the average CI minus the standard deviation was >1, a positive fitness effect was confirmed. Competition assays were performed in at least three independent experiments.

## 3 Results

### 3.1 Generation of a barcoded *S.* Choleraesuis TIS library and an ordered mutant collection

Our electroporations generated a *S.* Choleraesuis transposon mutant library consisting of 35,432 independent mutants with mapped Tn5 insertions, 34,811 of which were mapped to the genome and 621 to the plasmid. The mapped insertions disrupted 3,684 of 4,624 annotated chromosomal genes (80%), and 53 of 57 annotated plasmid genes (93%). Many of the undisrupted genes are probably essential for SCH under laboratory conditions. The library also contained insertions in 1,579 intergenic regions.

Additionally, we generated an ordered SCH transposon mutant collection consisting of 5,304 individual clones. This collection represents 2,838 of the 4,682 annotated SCH genes, corresponding to 61% coverage of the coding genome. For the 1,659 genes with multiple available mutants, two clones were included per gene, with priority given to insertions located on opposite strands and positioned near the C- or N-terminus while avoiding the first or last 10 bases of the gene. In addition to coding regions, the library also includes mutants affecting 807 of the 3,807 annotated intergenic regions. Overall, this curated library provides a broad representation across the SCH genome, enabling systematic functional analyses of both genes and intergenic elements.

### 3.2 *S.* Choleraesuis and native microbiota dynamics on alfalfa sprouts

Statistical analysis of the barcoded SCH TIS library’s dynamics on alfalfa sprouts revealed no statistically significant differences compared to the population dynamics of its WT (*p*_adj_>0.05, one-way ANOVA) for any of the sampled time points **(Figure S1)**.

Uninoculated alfalfa sprout samples were prepared and incubated under the same conditions as the screening samples to monitor the initial background microbiota and its dynamics in the absence of *Salmonella*. There was a high microbial load on alfalfa sprouts with initial MAB counts reaching 9.4 log_10_ CFU/g and *Enterobacteriaceae* populations approaching 8.0 log_10_ CFU/g. The bacterial load remained stable during the full incubation period (96 h) at 8°C **(Figure 1; Panel A)** with low variability among samples. No macroscopic changes were observed in the appearance of the inoculated alfalfa sprouts during the incubation period, consistent with commercially available sprouts.

**Figure 1.**
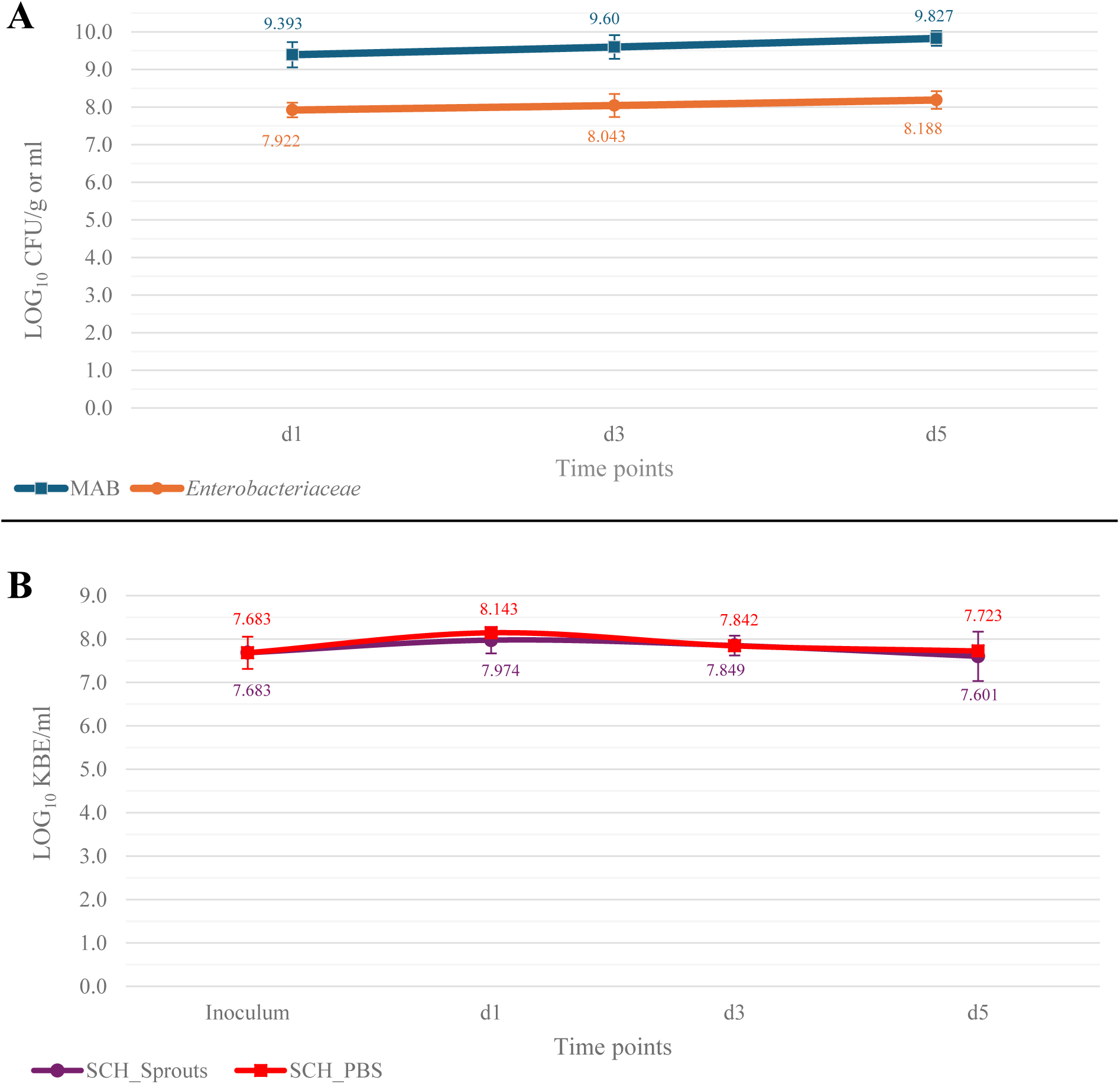
Population dynamics on alfalfa sprouts at 8°C. Panel. **A** illustrates the dynamics of the background microbiota, represented by mesophilic aerobic bacteria (MAB) and *Enterobacteriaceae*, at 1h (d_1_), 48h (d_3_), and 96h (d_5_) post-inoculation. **Panel B** shows the quantitative assessment of the *S.* Choleraesuis (SCH) barcoded TIS library in the inoculum and at the same sampling time points on alfalfa sprouts (SCH_Sprouts) and in PBS (SCH_PBS). There was no significant (*p* > 0.05) difference between the SCH library populations on alfalfa sprouts and in PBS during cold storage. Data represents averages from five biological replicates, with error bars indicating standard deviations. SCH: barcoded TIS library in *S.* Choleraesuis 11-Ga-2015.

The dynamics of the SCH library on alfalfa sprouts and in PBS at 8°C were not significantly different (*p*_adj_>0.05) and remained mostly stable during the incubation period **(Figure 1; Panel B)**.

### 3.3 Identification of *S.* Choleraesuis genes with a role during cold storage on alfalfa sprouts

Using our stringent two-pronged threshold for significance, only eight SCH genetic elements showed a statistically significant fitness effect during a 5-day cold storage on alfalfa sprouts when analyzed with the SCH random transposon library: six coding genes (*eda, fabF, lpp1_2, pnp, stpA, SCHChr_03621;* **Table 1**) and two intergenic regions (IRs) located between *deaD* and *nlpI* as well as between *pipB* and *sigE.* Further analyses of the genes delimiting the IRs showed that, while *nlpI* mutants remained mainly stable during the interaction with alfalfa sprouts, *deaD* mutants were negatively selected with stronger reductions in the sprout samples than in the negative controls but did not meet the statistical threshold for the sprouts versus *no matrix* comparison. While the insertion within the *pipB-sigE* IR was deleterious, disruption of the two flanking genes caused no fitness effects. Mutant dynamics during the screening period are represented in **Figure 2**.

**Figure 2.**
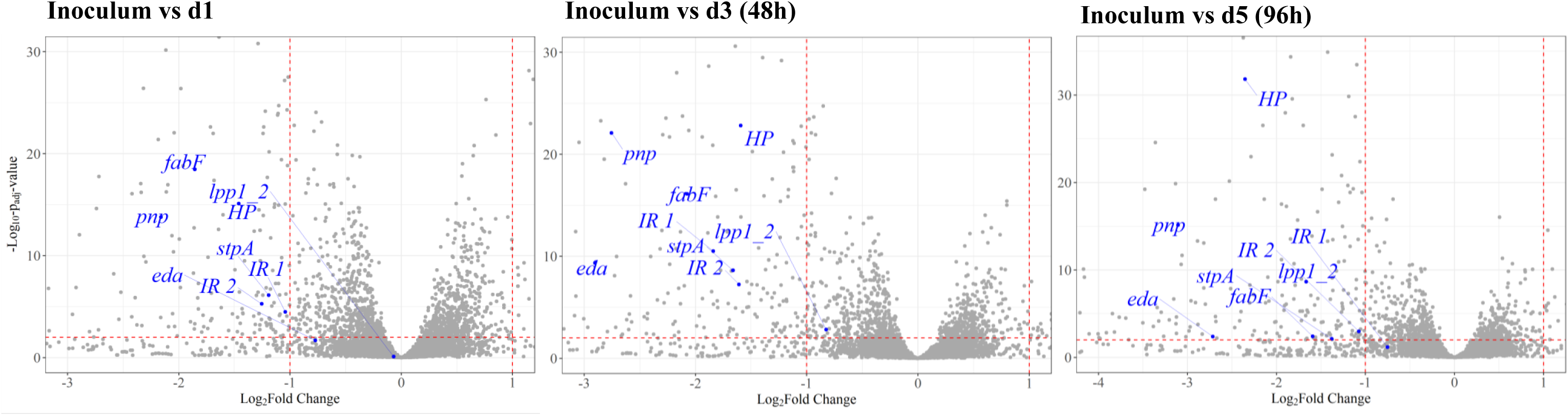
Changes in mutant abundances of *S.* Choleraesuis 11-Ga-2015 on alfalfa sprouts at 8°C. Volcano plots illustrate genome-wide fitness effects, with the log_2_ fold-change in relative mutant abundance on the X-axis and the adjusted *p*-values on the Y-axis. Comparisons are shown between the inoculum and samples collected 1h post-inoculation (d_1_), 48h post-inoculation (d_3_), and 96h post-inoculation (d_5_). Each dot represents an individual insertion mutant, and mutants which fulfilled the dual significance criterion are labelled in blue. Red dotted lines indicate the thresholds for significance in each comparison.

**Table 1.**
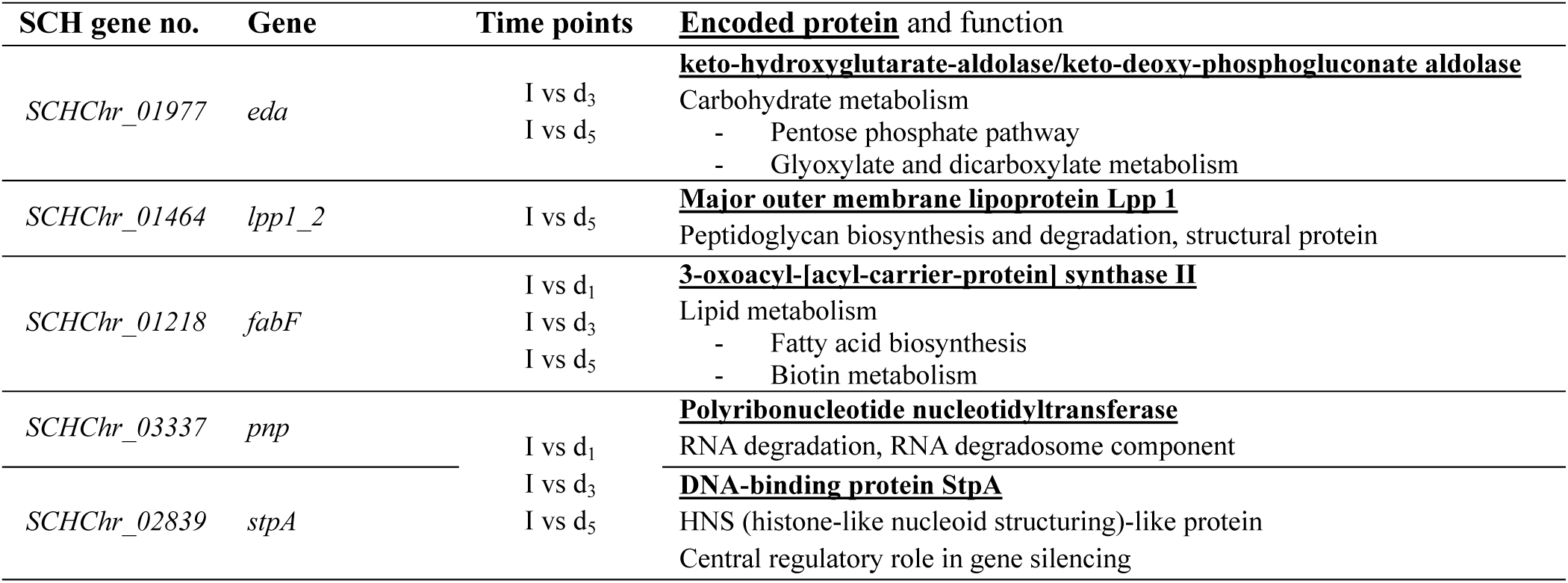
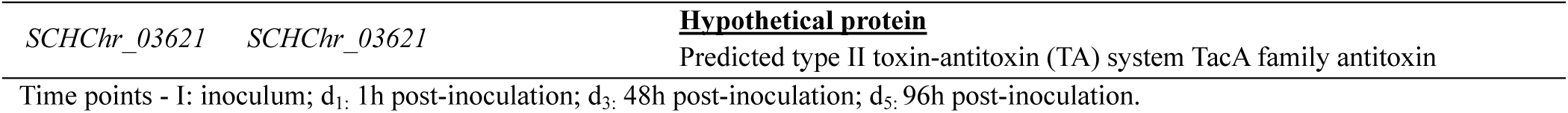
Genes with insertions resulting in a significant fitness effect in *S.* Choleraesuis on alfalfa sprouts in at least one comparison between time points.

KEGG enrichment analysis of genes affecting the fitness of SCH during cold storage on alfalfa sprouts was performed to provide an overview of relevant pathways for the interaction of *S.* Choleraesuis with this matrix. “Flagellar assembly” was significantly enriched (*q*≤0.01) together with three associated GO terms (*p*≤0.01), as summarized in **Table 2**. Analysis of the transposon data revealed that all selected genes associated with “flagellar assembly” exhibited an overall positive selection, which was particularly pronounced during the first hour of interaction but shifted to a mild negative effect between 1h post-inoculation and the final time point (96 h).

**Table 2.** Significantly enriched KEGG pathway and associated GO terms for the interaction of *S.* Choleraesuis on alfalfa sprouts at 8°C.

| KEGG Pathway | q-value | Selected Genes | GO IDs | GO Term | Classic Fisher |
| --- | --- | --- | --- | --- | --- |
| Cellular Processes<br>Cell motility<br>Flagellar assembly | 4.72 E-04 | <u>*flgAE*</u>, <u>*flhC*</u>,<br><u>*fliGK*</u>, <u>*fljB*</u> | GO:0051674 | Localization of the cell | 8.6 E-04 |
|  |  |  | GO:0071973 | Bacterial-type flagellum-dependent cell motility | 1.0 E-03 |
|  |  |  | GO:0040011 | Locomotion | 8.6 E-03 |

Listed genes fulfilled $p_{\text{adj}} < 0.05$ in at least one alfalfa sprouts/negative control and inoculum/timepoint comparison.

Additionally, GO terms “nitrogen fixation” (GO:0009399; *glnGA*) and “enterobacterial common antigen biosynthetic process” (GO:0009246; *wecBE, wzyE, rffM*) were significantly enriched.

### 3.4 Competition assays confirm genetic determinants

Insertion mutants in *eda, fabF, lpp1_2, mnmG,* and *stpA* were selected from the available mutants in the ordered SCH library collection for direct competition against the WT **(Table 3)**. Among these, *mnmG* (also named *gidA*) did not meet the dual requirement for fitness effect significance but was included due to the strong reduction in the mutant abundances and previous TIS data suggesting a recurring pattern on this food matrix (Führer et al., 2025).

**Table 3.**
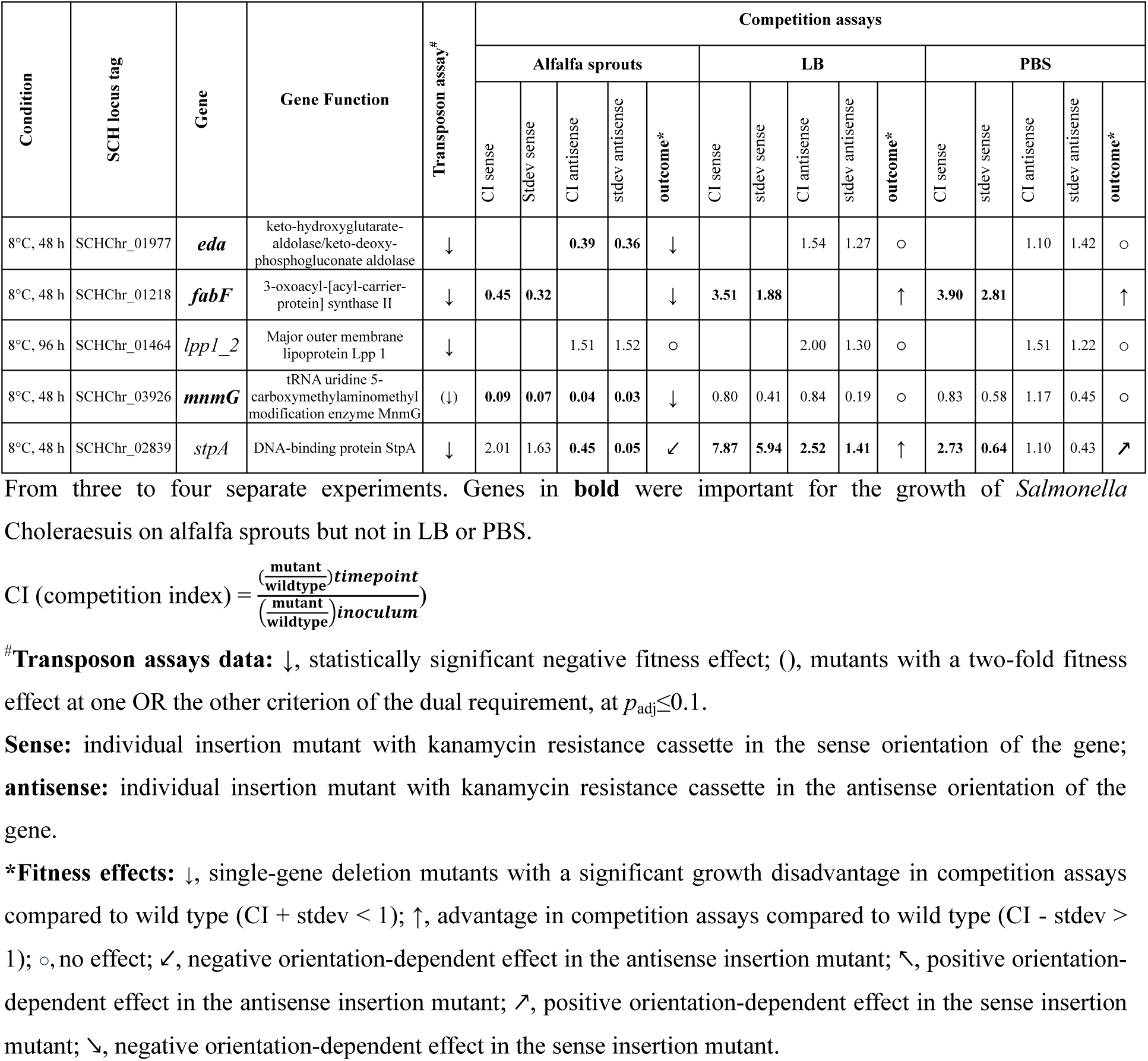
Fitness effect of selected *S.* Choleraesuis 11-Ga-2015 individual insertion mutants in competition with the wild type strain on alfalfa sprouts.

For *stpA* and *mnmG,* two individual IMs were available in both the sense and antisense orientations. For the remaining three genes, only one mutant was present – *fabF* with the transposon inserted in the sense orientation, and *eda* and *lpp1_2* with insertions in the antisense orientation. The sampling time point selected for the competition assays corresponded to that showing the strongest fitness effect in the transposon screen. Assays were performed on alfalfa sprouts and in nutrient-rich LB broth as a control to distinguish fitness effects associated with general nutrient availability from those specific to the alfalfa sprout environment. Competition assays were also run in PBS control conditions. The results are summarized in **Table 3**.

A negative fitness effect was confirmed when the competition index (CI) plus the standard deviation (STDEV) was less than 1 for both IMs, while a positive fitness effect was defined by a CI – STDEV > 1 for both mutants. For genes with only one available IM, meeting the described threshold in that single mutant was considered sufficient to validate the phenotype observed in the transposon assay.

## 4 Discussion

This study investigated fitness determinants of the plant-derived *S. enterica* serovar Choleraesuis strain 11-Ga-2015 on alfalfa sprouts during cold storage at 8°C. Quantitative analysis showed that overall *S.* Choleraesuis TIS library populations increased slightly within the first hour of interaction with sprouts but subsequently declined over the remaining time of storage. Although data on *S. enterica* growth dynamics on sprouts under cold conditions remain limited, our previous analyses of *S.* Typhimurium and *S.* Enteritidis on alfalfa sprouts under identical experimental conditions (Führer et al., 2025) showed no initial growth phase.

The consistently high background microbiota observed on alfalfa sprouts, as reported here and in previous studies (Führer et al., 2025; Jang et al., 2021; Young Kim et al., 2022), together with their association with multiple foodborne outbreaks (Miyahira & Antunes, 2021) – most commonly involving *Salmonella* and enterohemorrhagic *E. coli,* highlights the potential of sprouts to serve as a favorable niche for microbial persistence and growth. This aligns with the outcomes of our transposon screens of *S. enterica* libraries on alfalfa sprouts, where only a limited number of genetic elements showed significant fitness effects, what could be associated with the relatively permissive conditions of alfalfa sprouts at 8°C. Specifically, mutations in six *S. Choleraesuis* genes and two IRs conferred a fitness disadvantage, whereas 13 genes and five elements were identified in *S.* Enteritidis and *S. Typhimurium*, respectively, in our previous study (Führer et al., 2025).

Mutants carrying insertions in genes associated with carbohydrate metabolism (*eda*), RNA degradation (*pnp*), lipid metabolism (*fabF*), cell wall structure and interaction mechanisms (*lpp1_2*), DNA structuring (*stpA*), and a hypothetical protein (*SCHChr_03621*) exhibited significant fitness effects under the tested conditions. Competition experiments in *eda, fabF, lpp1_2, stpA,* and *mnmG* SCH mutants confirmed negative fitness effects for *eda, fabF,* and *mnmG* specifically in alfalfa sprouts, but not in nutrient media or PBS. These findings may underscore the importance of carbohydrate and lipid metabolism, as well as RNA degradation, for fitness of SCH in this ecological niche.

The pronounced fitness defect observed in *eda* mutants highlights the critical role of the carbohydrate metabolism – and specifically the Entner-Doudoroff (ED) pathway – in SCH fitness on alfalfa sprouts. The *eda* gene encodes KDPG (2-keto-3-deoxy-6-phosphogluconate) aldolase, a central enzyme of the ED pathway that operates in parallel with the classical Embden–Meyerhof–Parnas (EMP) glycolysis. Although the ED pathway yields only half as much ATP as the EMP pathway, recent work has revealed that it confers a distinct selective advantage under dynamically changing nutrient conditions (Law et al., 2024). Specifically, the ED pathway enables rapid metabolic acceleration upon carbon and nitrogen upshifts. This capacity for rapid flux adaptation likely provides a competitive advantage in fluctuating microenvironments such as plant surfaces, where nutrient availability is intermittent. Consistent with this, the ED pathway was shown to be required in the challenging environment inherent to various host infection processes, promoting *Salmonella* proliferation within macrophages and epithelial cells (Eriksson et al., 2003; Mitosch et al., 2023).

The relevance of this pathway is further underscored by the fitness effects of *S. enterica eda* mutants on various foods of plant origin. On melons, *eda* was critical for both *S.* Typhimurium and *S.* Enteritidis at room temperature, whereas its contribution was marginal at 8°C (Esteban-Cuesta et al., 2026). The phenotype observed at ambient temperature likely reflects a rapid upshift in glucose availability when interacting with ready-to-eat melons, where cutting increases nutrient release, favoring the ED pathway’s capacity for fast metabolic engagement. On sprouts, *S.* Typhimurium *eda* mutants also exhibited a negative fitness effect at 8°C (Führer et al., 2025). Collectively, these observations highlight the importance for SCH, and more broadly, for *S. enterica,* of efficiently managing energy and carbon resources in various fresh produce matrices, such as sprouts and melon.

The effects observed for *pnp* mutants highlight the role of RNA degradation in bacterial adaptation to the plant-associated environment by contributing to resource recycling and optimization. The *pnp* gene encodes the enzyme polynucleotide phosphorylase (PNPase), a highly conserved component of the RNA degradosome with established roles in mRNA turnover, tRNA processing, and sRNA-mediated regulation (Clements et al., 2002). Compared to SCH, *pnp* mutants of *S.* Typhimurium or *S.* Enteritidis exhibited relatively mild fitness defects on sprouts (Führer et al., 2025). Nevertheless, the importance of *pnp* for environmental resilience is evident across other produce-associated contexts: it contributes to the survival of *S.* Typhimurium on ready-to-eat muskmelons (Esteban-Cuesta et al., 2026) and onions (Führer et al., 2025), and transposon analyses similarly implicate it for *S.* Enteritidis and *S.* Newport fitness on muskmelons (Esteban-Cuesta et al., 2026). Disruption of *pnp* also reduces fitness on low-moisture foods, including almonds (*S.* Enteritidis) and pistachios (*S.* Typhimurium) (Li et al., 2020). Competition assays in nutrient-rich conditions (LB) provided an important control: while *S.* Typhimurium *pnp* mutants showed no fitness defects at 22°C (Esteban-Cuesta et al., 2026), they exhibited a significant disadvantage at 8°C on onions (Führer et al., 2025) and in this study, consistent with the established role of PNPase in the bacterial cold-shock response. Although *S. enterica pnp* mutants display milder cold sensitivity compared to other bacteria (Clements et al., 2002), their reduced fitness at low temperatures underscores the enzyme’s importance for fitness under cold stress.

We observed a significant fitness effect in SCH *fabF* mutants on alfalfa sprouts. FabF (β-ketoacyl-acyl carrier protein synthase II) catalyzes fatty acid (FA) elongation, particularly contributing to the synthesis of long-chain and unsaturated FAs (Guest et al., 2021). This activity is critical for maintaining membrane fluidity, permeability, and overall cell envelope integrity. Although SCH encodes other β-ketoacyl-ACP synthases, including FabB, these enzymes cannot fully compensate *fabF* mutations: in addition to catalyzing FA elongation, FabF also modulates chain length in response to temperature changes and is less sensitive to inhibitors like cerulenin (Guest et al., 2021). This further highlights the specialized role of FabF in envelope maintenance.

The observed fitness defect in SCH *fabF* mutants was specific to alfalfa sprouts and absent in LB or PBS, indicating that the contribution of FabF is food-matrix dependent and particularly critical for SCH survival on alfalfa sprouts under cold storage. Low temperatures reduce membrane fluidity, and bacteria adapt by increasing the proportion of unsaturated fatty acids to maintain membrane homeostasis (Ricke et al., 2018). Consistent with this role, *fabF* mutants in *E. coli* (Garwin et al., 1980), *Clostridium acetobutylicum* (Zhu et al., 2009), and deep-sea bacteria (Allen & Bartlett, 2000) exhibit cold-sensitive phenotypes, highlighting the conserved function of FabF in cold adaptation.

Membrane lipid composition not only changes in response to cold stress, but also as a result of desiccation. Accordingly, *fabF* mutants exhibited a negative fitness effect under desiccation stress in *S.* Typhimurium on pistachios stored at 25°C (Jayeola et al., 2020), and in *E. coli*, desiccation triggers adjustments in FA composition to preserve membrane fluidity (Scherber et al., 2009). Together, these observations underscore the central role of FA metabolism in maintaining membrane integrity.

The observed fitness effects of *fabF* and *lpp* mutants are likely interconnected. In *Salmonella,* the outer membrane murein lipoprotein Lpp is encoded by two genes, *lpp1* and *lpp2*, and disruption of either results in reduced virulence strains (Sha et al., 2004). Proper membrane function depends on maintaining bilayer integrity, which is supported in part by the regulation of lipid A biosynthesis (Guest et al., 2021). Deficiencies in FA biosynthesis mediated by *fabF* can indirectly perturb the metabolism of envelope-associated polysaccharides, including lipopolysaccharides (LPS) and the enterobacterial common antigen (ECA), because its products supply the phospholipids for the inner leaflet of the outer membrane (Guest et al., 2021; Zhou et al., 2025). LPS, including genes involved in O-antigen biosynthesis, have previously been shown to play an important role in the interaction of *S. enterica* with sprouts (Führer et al., 2025; Holden et al., 2024) as well as with other foods of plant origin, including low-moisture foods, onions, and melons (Esteban-Cuesta et al., 2026; Führer et al., 2025; Jayeola et al., 2020; Li et al., 2020).

This effect may also relate to the association of Lpp with the Tol-Pal system, a five-component transmembrane complex whose disruption compromises outer membrane integrity, alters LPS composition, and increases susceptibility to multiple stressors, including antimicrobial compounds such as polymyxins (Szczepaniak & Webby, 2024). Genome analysis of SCH showed that *lpp1* is represented by two CDS, *lpp1_1* and *lpp1_2.* Only mutants in *lpp1_2* were present in the SCH barcoded transposon library, and these exhibited a significant negative fitness effect on sprouts, whereas mutants in *lpp2 –* located between *lpp1_2* and *lpp1_1* in the antisense orientation – did not.

Competition assays between the *lpp1_2* SCH insertion mutant and the WT revealed no fitness disadvantage for the mutant under any of the tested conditions (sprouts, LB, and PBS). This may indicate functional redundancy between *lpp1_1* and *lpp1_2*, warranting further investigation through the construction of an *lpp1_1/2* double mutant. However, the apparent signal might also represent a bioinformatics artefact rather than a genuine biological phenomenon.

SCH mutants in *mnmG* (also referred to as *gidA*) exhibited pronounced reductions in fitness on alfalfa sprouts and, in previous studies, for *S.* Typhimurium on onions and sprouts (Führer et al., 2025). Although these effects fell just below the threshold of statistical significance relative to control samples, *mnmG* emerged as a gene of interest for the competition experiments, which confirmed a matrix-specific fitness defect for *mnmG* mutants on alfalfa sprouts.

MnmG (glucose-inhibited division protein A) is a highly conserved tRNA modification enzyme that, together with MnmE, catalyzes the addition of a 5- carboxylmethylaminomethyl group to the wobble uridine of tRNAs. This modification is critical for efficient and accurate translation, influencing global protein synthesis and regulation of diverse cellular processes. In *S. enterica*, *mnmG* also modulates the expression of virulence-associated genes, regulating pathogenesis *in vivo* and *in vitro* infection models, and influences growth, stress tolerance, and biofilm formation (Shippy & Fadl, 2014; Shippy et al., 2012).

Although a general fitness defect might be anticipated for *mnmG* mutants, the phenotype was particularly pronounced under the specific stress conditions present on alfalfa sprouts. This environment likely imposes a combination of selective pressures, such as variable nutrient availability, plant-derived oxidative and antimicrobial compounds, and challenges associated with colonization and cold storage conditions, which together may contribute to the critical role of *mnmG* in SCH fitness in this ecological niche.

The *SCHChr_03621* mutant showed a significant fitness defect in SCH in the transposon assay. Sequence comparison using the Uniprot database (Ahmad et al., 2025) and blast+ (Camacho et al., 2009) predicts that *SCHChr_03621* encodes the antitoxin component of a TacA/TacT family type II toxin-antitoxin (TA) system. Non-secreted type II TA system toxins in *Salmonella* are induced inside host macrophages, leading to growth arrest and the formation of metabolically active, non-growing persister populations (Helaine & Kugelberg, 2014; Maisonneuve & Gerdes, 2014). TacT induces growth arrest in *S.* Typhimurium by stalling the ribosome, a process that is reversible by the activity of the antitoxin TacA (encoded by *pth*) (Cheverton et al., 2016). We therefore hypothesize that conditions on alfalfa sprouts may similarly trigger TacT expression, rendering the loss of the antitoxin *tacA* detrimental to bacterial fitness.

Competition experiments between SCH *stpA* mutants and the WT revealed an orientation-dependent negative fitness effect in the antisense orientation of *stpA.* Located upstream of *stpA* is *SCHChr_02838,* a rhodanese-related sulfurtransferase encoded in a separate operon. Insertion mutants in this locus did not display significant fitness defects in the transposon assays on alfalfa sprouts, indicating the fitness defect is unrelated to *SCHChr_02838*. Examination of the transposon insertion sites of both *stpA* insertion mutants revealed that the sense-oriented insertion is positioned near the 3’ end of the gene, whereas the antisense insertion lies centrally within the coding sequence. This positional difference could indicate that the sense-oriented insertion may not disrupt essential functional domains of StpA, suggesting the observed fitness defect would correspond to the disruption of *stpA* itself.

Flagellar assembly was significantly enriched in both KEGG and GO analyses, with mutant abundances of six selected genes showing an initial negative selection that shifted to positive over time. This dynamic has been observed in previous studies on alfalfa sprouts (Führer et al., 2025; Holden et al., 2024) and melons (Esteban-Cuesta et al., 2026). We hypothesize that the loss of flagella may initially disadvantage individuals by reducing their ability to seek nutrient-rich niches and avoid stressors. However, the plant immune system can detect bacterial flagella, and mutants may gain a competitive advantage during colonization, as was shown in *Listeria monocytogenes* (Gorski et al., 2009). The loss of flagella during interactions with plant matrices may represent a conserved bacterial strategy to evade plant immune defenses; however, it may also reflect an energetic trade-off, as flagella are of limited functional benefit in our experimental system and energetically costly under resource-limited conditions. Previous studies have suggested that other attachment structures, such as aggregative fimbriae and fimbrial regulatory systems, may play a more relevant role in the attachment of *Salmonella* to plants (Barak et al., 2005; Holden et al., 2024). However, competition experiments using *flhDC* and *flhA* deletion mutants revealed reduced ability of these mutants to attach to sprouts, contrary to their TraDIS-Xpress results (Holden et al., 2024).

Overall, our study identifies key genetic and functional traits that contribute to *S. enterica* fitness on alfalfa sprouts during cold storage, thereby extending insights beyond the traditionally studied serovars by focusing on a strain that is capable of internalization within plant tissues (Esteban-Cuesta et al., 2018). Although this serovar is typically host-adapted to pigs, its ability to colonize plants provides a valuable model for comparing specific mechanisms and adaptations across different hosts and environmental niches. The identified genomic factors highly overlap with known determinants of plant–microbe interactions and bacterial competition, including oxidative stress mitigation, cell envelope structural components, and motility-associated traits, highlighting the shared strategies employed by enteric pathogens to persist and compete within plant-associated environments.

## 5 Acknowledgements

We would like to thank Larissa Klose, Julia Völker, Johanna Dietz, and Sandra Haid for their excellent technical assistance during the competition experiments, as well as Julia Mühmel and Meike Schumann for their support in preparing laboratory media. We greatly appreciate their expertise and contributions.

This project is supported by a grant awarded to Dr. Irene Esteban Cuesta by the German Research Foundation (Deutsche Forschungsgemeinschaft, DFG) with Project No. 494875429.

## 6 Declaration of generative AI and AI-assisted technologies in the writing process

During the preparation of this manuscript, the authors used ChatGPT-5 to enhance readability and clarity. All content generated with these tools was thoroughly reviewed and edited by the authors, who take full responsibility for the final version of the manuscript.

## 7 Supporting Information

Supplementary Table S1. Summary of the primers used in the study.

Supplementary Table S2. DESeq2 statistical analysis for the transposon insertion sequencing experiments of the barcoded *S.* Choleraesuis library on alfalfa sprouts.

Supplementary Figure S1. Comparison of the population dynamics of the *S.* Choleraesuis 11-Ga-2015 library and the wild type strain.

